# Conservation of locomotion-induced oculomotor activity through evolution in higher tetrapods

**DOI:** 10.1101/2021.06.26.450039

**Authors:** Filipa França de Barros, Julien Bacqué-Cazenave, Coralie Taillebuis, Gilles Courtand, Marin Manuel, Hélène Bras, Michele Tagliabue, Denis Combes, François M Lambert, Mathieu Beraneck

## Abstract

Efference copies are neural replicas of motor outputs used to anticipate the sensory consequences of a self-generated motor action or to coordinate neural networks involved in distinct motor behaviors^1^. An established example of this motor-to-motor coupling is the efference copy of the propulsive motor command that supplements classical visuo-vestibular reflexes to ensure gaze stabilization during amphibian larval locomotion^2^. Such feedforward replica from spinal pattern-generating circuits produces a spino-extraocular motor coupled activity that evokes eye movements, spatio-temporally coordinated to tail undulation independently of any sensory signal^3,4^. Exploiting the evolutionary-development characteristic of the frog^1^, studies in metamorphing Xenopus demonstrated the persistence of this spino-extraocular motor command in adults, and its developmental adaptation to tetrapodal locomotion^5,6^. Here, we demonstrate for the first time the existence of a comparable locomotor-to-ocular motor coupling in the mouse. In neonates, *ex vivo* nerve recordings from brainstem-spinal cord preparation reveals a spino-extraocular motor coupled activity similar to the one described in Xenopus. In adult mice, trans-synaptic rabies injection in lateral rectus eye muscle labels cervical spinal cord neurons projecting directly to abducens motor neurons. Finally, treadmill-elicited locomotion in decerebrated preparations^7^ evokes rhythmic eye movements in synchrony with the limb gait pattern. Overall, our data are evidence for the conservation of locomotor-induced eye movements in higher tetrapods. Thus, in mammals as in amphibians, during locomotion CPG-efference copy feedforward signals might interact with sensory feedback to ensure efficient gaze control.

**Highlights:**

- Spino-extraocular motor coupling is evidenced from newborn mice ex vivo preparations
- Adult decerebrated mice exhibit conjugated rhythmic eye movements during treadmill locomotion
- Locomotor-induced oculomotor activity occurs in absence of visuo-vestibular inputs
- Conserved CPG-based efference copy signal in vertebrates with common features.

**eTOC blurb:** We report a functional coupling between spinal locomotor and oculomotor networks in the mouse, similar to the one previously described in Amphibians. This is the first evidence for the direct contribution of locomotor networks to gaze control in mammals, suggesting a conservation of the spino-extraocular coupling in higher tetrapods during sustained locomotion.

## Results

### Fictive locomotor activity induces ex vivo spino-extraocular motor coupling in neonates in absence of sensory inputs

The ability of locomotor CPG rhythmic activity to elicit spino-extraocular motor coupling was evaluated on isolated *ex vivo* brainstem-spinal cord preparations of neonatal mice (Figure 1A). Electrical stimulation of either the first sacral dorsal root (S1Dr, Figure 1A) or the 8th cervical dorsal root (Dr. C8, Figure S1A) evoked episodes of fictive locomotion with the typical coordination pattern between cervical and lumbar ventral roots (Vr) rhythmic activity (Vr; Figure 1A), as previously reported^8,9^. In this locomotor pattern, paired bursting discharge in homolateral C8 (light green in Figure 1A, B bottom panel) and L2 (light orange in Figure 1A, B bottom panel) Vr was in alternance with homolateral C5 and L5 Vr^9^. In most cases (66%, Figure 1C) this rhythmic cervico-lumbar coordinated locomotor activity was coupled to a bursting activity of the abducens motor nerves (Figure 1A-B) within the same frequency range (0.5-1.5Hz; Figure1D, E) and a latency of ∼62ms to C8 Vr burst and ∼80ms to L2 Vr burst, respectively (Figure 1F), compatible with a monosynaptic delay^10,11^. Inversely, non-coordinated spinal activity failed to evoke a coupled rhythmic discharge in abducens nerves (Figure 1C), which instead displayed a tonic, non-rhythmic activation. Locomotor-related bursts in the bilateral abducens occurred in alternation (Figure 1B, top panel) and were in synchrony with the ipsilateral paired C8-L2 Vr rhythmic discharge, respectively (Figure 1G). Frequency analysis of abducens discharge in response to S1Dr stimulation (Figure S1C-E) demonstrated that the spino-extraocular coupling was not induced by proprioceptive feedback inputs. In addition, coordinated fictive locomotor episodes evoked by NMDA/5HT spinal application also produced a well correlated rhythmic activity in the abducens motor nerves (Figure 1H - I) in a lower frequency range (0.15-0.25Hz; Figure 1J) and with a preserved phase relationship relative to the Vr burst discharge (Figure 1K). Since all these brainstem-spinal preparations lacked visuo-vestibular sensory inputs as well as cortical, mesencephalic and cerebellar inputs (see methods), this spino-extraocular motor coupling was produced by a locomotor efference copy signal, originating from the spinal motor CPGs.

**Figure 1.**
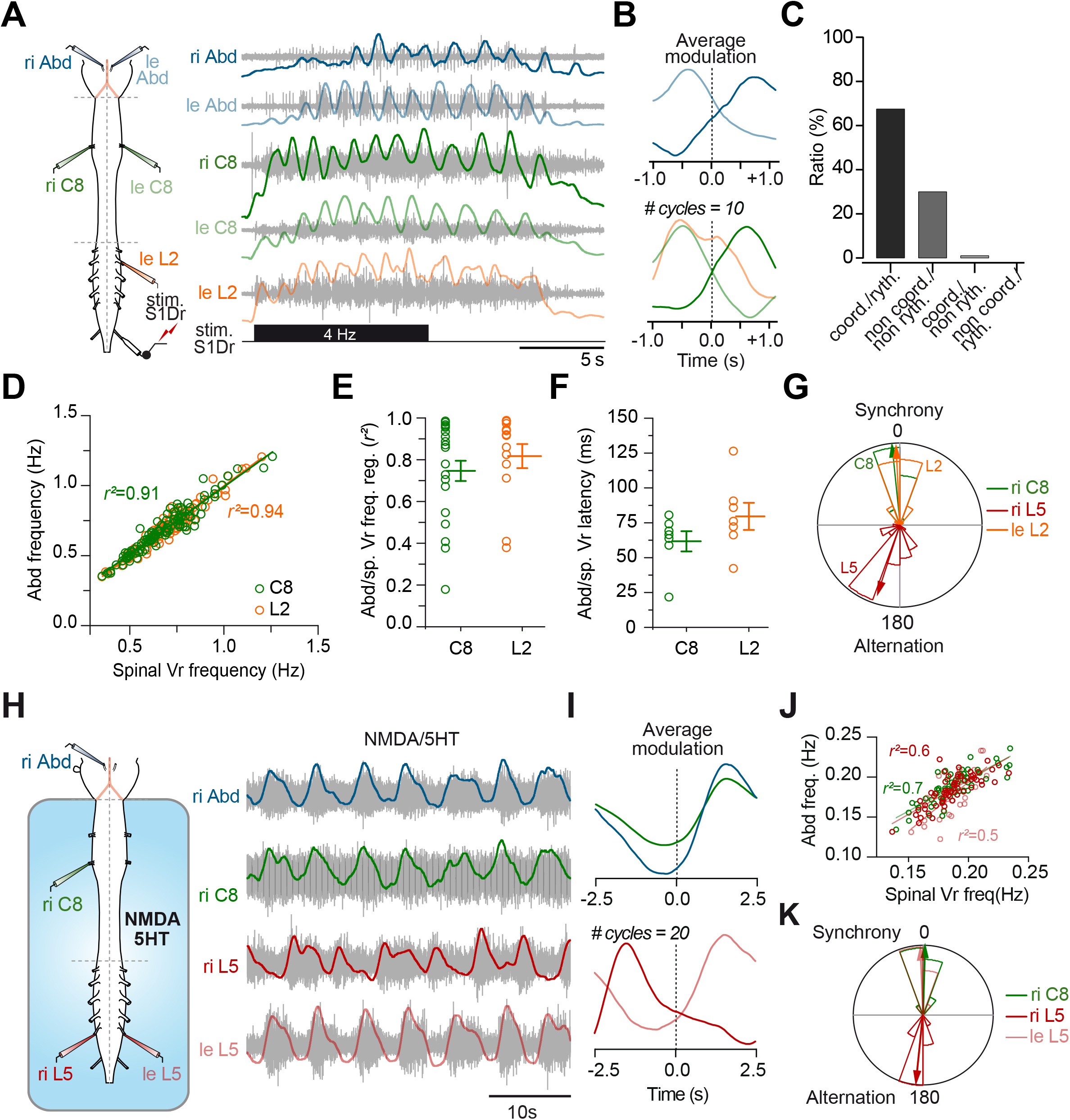
Fictive locomotor activity evokes spino-extraocular motor coupling in neonatal brainstem-spinal cord isolated preparations. **(A)** Schematic of the neonatal mouse brainstem and spinal cord preparation with stimulated and recorded nerve branches (left side). Extracellular nerve recordings (right side, raw traces in light gray, integrated traces in color) of the left (le, light blue) and right (ri, dark blue) abducens nerves (Abd.), the left and right 8th cervical roots (le C8, light green; ri C8, dark green) and the right 5th lumbar ventral root (ri L5, red). Discharges recorded during an episode of fictive locomotion evoked by the electrical stimulation of the S1 dorsal root (stim. S1Dr) with a 4Hz pulse train (black rectangle). **(B)** Average cyclic modulation of the discharge activity (integrated trace) from the motor nerves shown in (A) over 10 consecutive locomotor cycles. The ri C8 trace (dark green) was used as the reference to determine locomotor cycles. **(C)** Percentage of total (n=7 mice) preparations with a coordinated fictive locomotor pattern coupled with a rhythmic abducens discharge (coord./ryth, black histogram; 66%), with an absence of coordinated fictive locomotor pattern and an absence of rhythmic abducens discharge (non coord./non ryth, dark gray histogram; 33%), with a coordinated fictive locomotor pattern and an absence of rhythmic abducens discharge (coord./non ryth, light gray histogram; 1%), with an absence of coordinated fictive locomotor pattern and a rhythmic Abducens discharge (non coord./ryth; 0%). **(D)** Linear correlation in bursting frequencies between the abducens (Abd) nerve discharge and the C8 (green) (Abd vs C8, n= 7, R = 0,9567, Pearson test, *p <0,0001*; = 0,9153), L2 (orange) (Abd vs L2, n = 7, R= 0,9696, Spearman test, *p <0,0001;* r² =0.9400) spinal ventral roots. **(E)** Individual (empty circles) and mean (bars) ±SEM of the linear regression (r²) and the **(F)** latency between firing of the abducens and spinal root (milliseconds, ms) in bursting frequencies (freq.) between the abducens (Abd) nerve discharge and C8 (green, mean r²= 0,75±0,06, mean latency = 61.7 ± 7.2ms), L2 (orange, mean r²= 0,82±0,06, mean latency= 79.6 ± 9.7ms) Sp. Vr. discharges for each preparation, independently of the frequency. **(G)** Circular plots showing the phase relationships between the discharge burst in the abducens nerve and ipsilateral C8 (µ = 354,288° ± 1,467; r= 0, 97, n= 7), L2 (µ = 357,359° ± 2,213; r = 0,936; n= 8) and L5 (µ= 198 ± 5,04; r= 0,868, n= 3) Sp. Vr. In this and all polar plots, the width of the wedges is 0,05. **(H)** Schematic of the neonatal mouse brainstem and spinal cord preparation with recorded nerve branches (left side). Extracellular nerve recordings (right side, raw traces in light gray, integrated traces in color) of the right (ri, dark blue) abducens nerves (Abd.), the right 8th cervical roots (ri C8, dark green) and the right and left 5th lumbar ventral root (ri L5, dark red; le L5, light red) discharges during an episode of fictive locomotion evoked by bath application of 5HT (15 µM/NMDA (7,5 µM) restricted to the spinal cord. **(I)** Averaged cyclic modulation of the discharge activity (integrated trace) of the motor nerves shown in panel H over 20 consecutive locomotor cycles. The ri C8 trace (green) was used as the reference to determine locomotor cycles. **(J)** Linear correlation in bursting frequencies between the abducens (Abd) nerve discharge and C8 (green) (Abd vs C8, n= 2, R = 0,8187, Spearman test, *p <0,0001*; r² = 0.6702), ipsilateral L5 (ri L5, dark red) (Abd vs L2, n= 2, R= 0,7879, Spearman test, *p <0,0001;* r² =0.6208) and, contralateral L5 (le L5, light red)(Abd vs L5, n= 2, R= 0,7045, Spearman test, *p <0,0001;* r² =0.4964) spinal ventral roots (Vr) discharges from the sequence shown in H. **(K)** Circular plots showing the phase relationships between the discharge burst in the abducens nerve and ipsilateral C8 (µ= 1,432°±2,52; r= 0,949), ipsilateral L5 (µ= 357,97° ±1,826; r=0,9712) and contralateral L5 (µ= 185,835°± 2,273; r= 0,954) spinal ventral roots (n=2 mice). The ri C8 and the ri L5 were in synchrony with the abducens nerve discharge while the L5 was out of phase.

### Spino-extraocular motor coupling depends on the integrity of the cervical, and not lumbar, spinal cord

To determine the respective roles of the cervical and lumbar CPG networks in the origin of locomotor-induced spino-extraocular motor coupling, cervical and lumbar regions were selectively isolated in brainstem-spinal cord preparations with spinal split-baths (Figure 2A). When the cervical cord was perfused with a calcium-free aCSF (Figure 2A, left) electrical stimulation of the S1Dr evoked typical alternated fictive locomotor activity in the lumbar Vr (Figure 2B bottom traces, Figure 2D upper panel). However, it failed to evoke correlated fictive locomotor activity in the cervical Vr or rhythmic discharge in the abducens motor nerves (Figure 2B upper traces, Figure 2D, lower panel). This experiment demonstrates the instrumental role of cervical motor networks activity in the genesis of the spino-extraocular motor coupling. Next, we tested whether the cervical cord acted as a neuronal relay or was the origin of the spino-extraocular motor coupling. Locomotor-induced rhythmic discharge of abducens motor nerves was recorded in response to electrical stimulation of either S1Dr or C8Dr, in isolated *ex vivo* preparations exposed to a mid-thoracic section (Figure 2E). After the thoracic section, S1Dr stimulation failed to produce any spino-extraocular motor coupling (Figure 2F) whereas C8Dr electrical stimulation evoked locomotor-induced rhythmic discharge of the abducens motor nerves (Figure 2G). Even though the cervical CPG-related abducens rhythmic activity was recorded at relatively low frequency (Figure 2H, 0,2-0,35Hz, orange circles), it remained in synchrony with C8Vr bursting discharge (Figure 2I and 2J). Overall, these results show that, while the activation of the lumbar CPG allowed for a more robust coupling, the locomotor-related activity in abducens motor nerves originated specifically from the cervical CPG.

**Figure 2.**
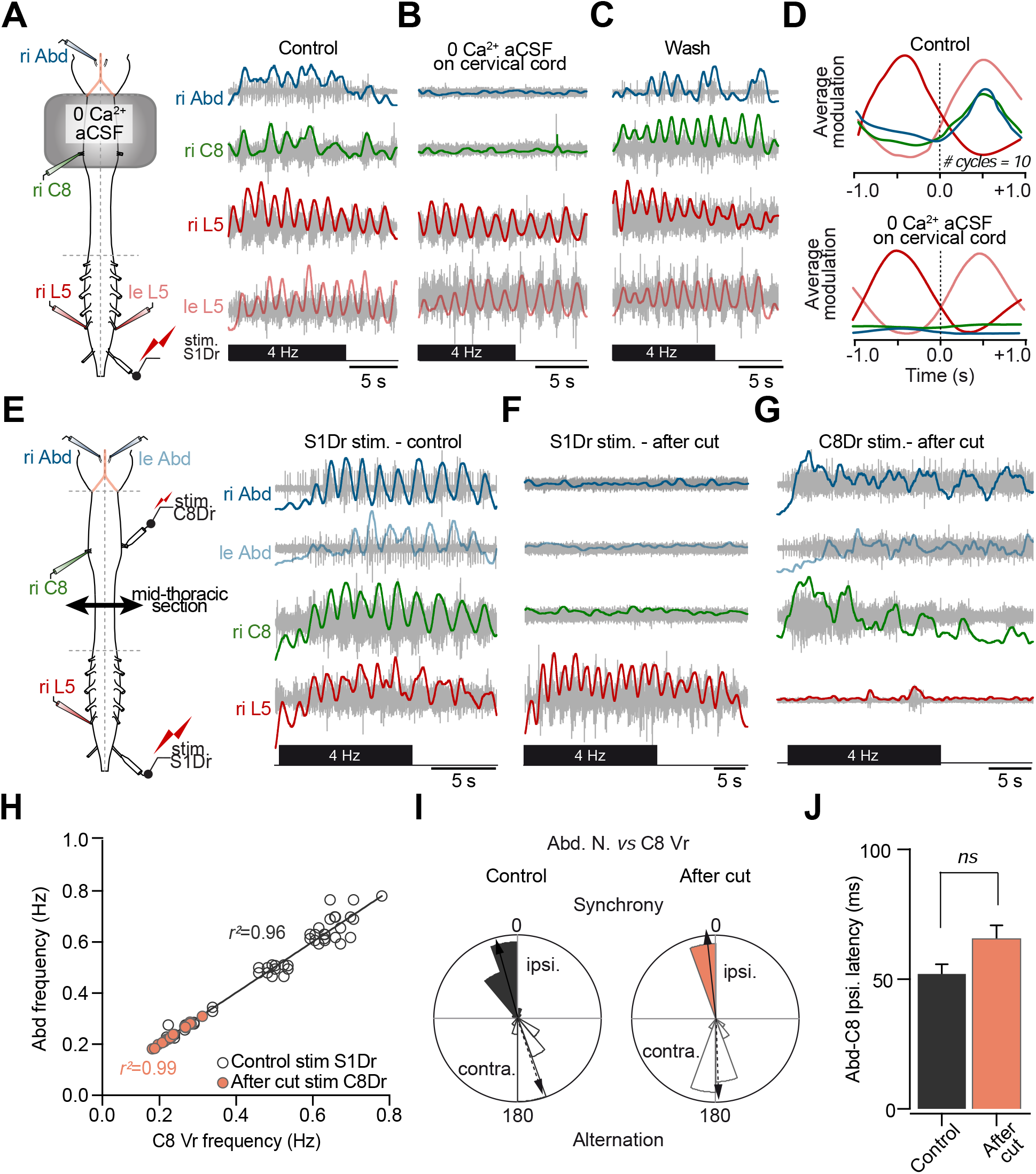
Efference copy signaling responsible for spino-extraocular motor coupling originates from cervical locomotor CPG. **(A)** Schematic of the neonatal mouse brainstem and spinal cord preparation with the recorded nerve branches (left side) during the calcium free (0Ca2+) aCSF experiments. Extracellular nerve recordings (raw traces in light gray, integrated traces in color) of the right (ri, dark blue) abducens nerves (Abd.), the right 8th cervical roots (ri C8, dark green) and the right and left 5th lumbar ventral roots (ri L5, dark red; le L5, light red) discharges during an episode of fictive locomotion evoked by the electrical stimulation of the S1 dorsal root (stim. S1Dr) with a 4Hz pulse train (black rectangle). In the control condition (**A**), during bath application of 0Ca2+ aCSF (**B**) restricted to the cervical spinal cord and after washout (**C**). **(D)** Averaged cyclic modulation of the discharge activity (integrated trace) from the motor nerves shown in A (top panel, control condition) and B (bottom, 0Ca^2+^) over 10 consecutive locomotor cycles. The ri L5 trace was used as the reference to determine locomotor cycles. **(E)** Schematic of the neonatal mouse brainstem and spinal cord preparation with the recorded nerve branches (left side) during the mid-thoracic section experiments. Extracellular nerve recordings (raw traces in light gray, integrated traces in color) of the right (ri, dark blue) and left (le, light blue) abducens nerves (Abd.), the right 8th cervical roots (ri C8, dark green) and the right 5th lumbar ventral root (ri L5, dark red) discharges during an episode of fictive locomotion evoked by the electrical stimulation of the S1 dorsal root (stim. S1Dr) with a 4Hz pulse train (black rectangle). Responses for control (**E**), after mid-thoracic section (**F**) and after mid-thoracic section during an episode of fictive locomotion evoked by the electrical stimulation of the C8 dorsal root (**G**, stim. C8Dr) with a 4Hz pulse train. **(H)** Linear correlation in bursting frequencies between Abducens (Abd) nerve discharge and ispi C8 Vr discharge in response to stimulation of the S1Dr during control conditions (gray circles, R=0,9817, Pearson test, p<0,0001, n= 3; r2=0,9637), and after the mid-thoracic lesion (orange dots) (R=0,9983, Pearson test, p<0,001, n= 3; r2=0,9966). **(I)** Polar plots of the phase relationships between the abducens and C8Vr before (control, left) abd C8 ipsi before (µ= 345,161°±2,546; r= 0,974) abd C8 contra before (µ= 162,803±4,285; r= 0,943) and after mid-thoracic lesion (after cut, right) abd C8 ipsi after cut (µ= 353,912±2,463; r= 0,999), abd C8 contra after cut (µ= 177,873±4,909; r= 0,939)(n=3 mice). The phase relationship is conserved after the cut. **(J)** Mean absolute latency time between the activity in the C8 Vr and the abducens nerve in control (ctrl) (52,96±4,51ms) and after cut (66,49±5,96ms). There was no significant difference in the delay observed before and after the mid-thoracic cut (t-test, p=0,1038, n=3).

### Last-order pre-motor neurons in cervical cord project to abducens motoneurons

To determine the cervical neuronal relay implicated in the spino-extraocular coupling (Figures 1 and 2), transsynaptic retrograde tracing of last-order premotor neuronal populations was performed by rabies virus (RV) injection in the lateral rectus muscle (LRM) of adult mice (Figure 3A). The monosynaptic labeling time window (55h after RV infection of motoneurons innervating LRM, Figure 3A and 3B) revealed several populations of last-order premotor neurons connected directly to RV^+^ abducens motoneurons (visualized by ChAT immunolabeling, Figure 3B). As expected from vestibulo-ocular reflex circuitry, RV^+^ neurons were found in the bilateral medial vestibular nuclei (MVN, VOR neurons, Figure 3C). Moreover, RV^+^ neurons were observed bilaterally throughout the cervical spinal cord (Figure 3D-F) in all preparations (n=3; Figure 3G-I). Average RV infection ratio was of 1 abducens motoneuron contacted by 6 cervical neurons (Figure 3J). RV^+^ cervical neurons were mainly located in the spinal ventral horn or around the central canal (Figure 3K). Notably, at this monosynaptic infection time, no RV^+^ neurons were observed caudally from the C8 segment. An extended infection time (70h) revealed spinal neuronal populations disynaptically connected with abducens motoneurons, scattered throughout the gray matter as well as beyond the cervical region (Figure S3B). This result suggests that inputs from spinal motor networks and sensory spinal circuitry converge at least on the direct abducens spino-extraocular pathway (Figure S3A). Overall, the use of RV trans-synaptic injections revealed a spinal-brainstem pathway likely to support the functional coupling of abducens motor nerve discharge with the rhythmic activity evoked by the cervical-CPG during locomotion in mice.

**Figure 3.**
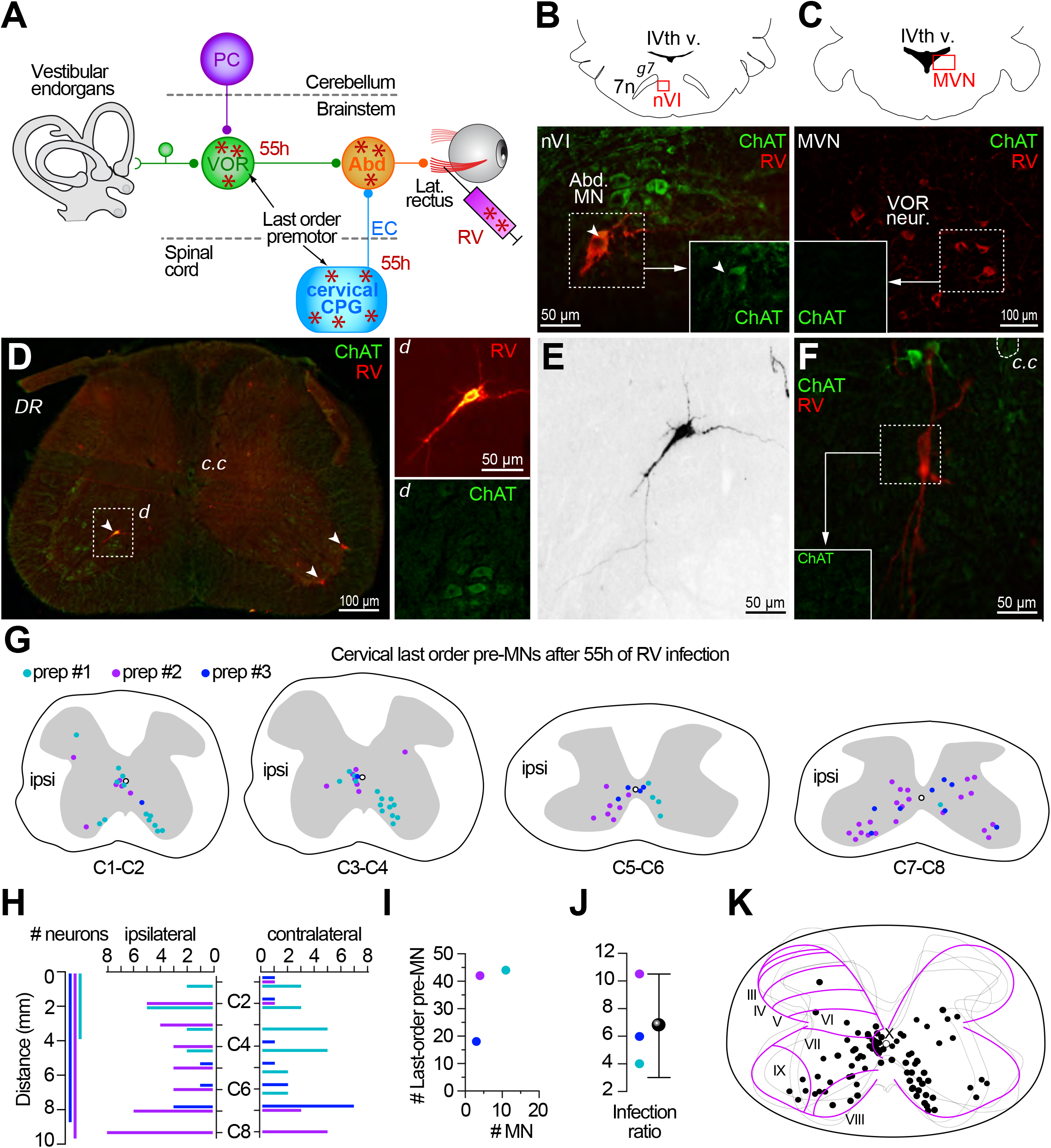
Anatomy of the spino-extraocular pathway. **(A)** Depiction of the rabies virus (RV) injection protocol. RV injections were performed in the lateral rectus muscles of the left eye. After 55h of RV migration, labeled neurons (red asterisks) were found in the abducens (Abd) nuclei, and in the mono synaptically connected last-order premotor structures. PC, Purkinje cells; VOR, Vestibulo-ocular central neurons; EC, efferent copy spinal neuron; cervical central pattern generator, cervical CPG. **(B)** Representation (top panel) of the location (red box) of the RV+ neurons in the abducens nucleus (nVI) and its landmarks: geniculum of the facial nerve (g7) and the 4th ventricle IVth v). The bottom panel shows a fluorescence microscopy image (20x) of an RV+/ChAT+ neuron (RV, red) and abducens motoneurons (AbD. MN, ChAT, green) 55h after the infection. **(C)** Representation (top panel) of the location (red box) of the RV+ neurons in the medial vestibular nucleus (MVN) 55h after the infection. The fluorescence microscopy image (bottom, 20x) shows RV+ VOR neurons in the MVN. **(D)** Fluorescence microscopy image (5x magnification) of a cervical spinal cord slice and an example location of RV+ neurons in the ventral horn ((d) top image, white arrowheads) as well as ChAT+ neurons ((d) bottom image). C.c, central canal; DR, dorsal root. **(E)** Black and white contrasted image of the neuron shown in inset *d*,D showing its neurite ramifications (20x magnification, scale bar = 50µm). **(F)** Example of a RV+ last-order premotor neuron located near the central canal (c.c). No ChAT staining was observed (ChAT, empty box), excluding the possibility of the shown neuron being a motoneuron. The dendrites of the RV+ are in close proximity with the ChAT+ neurons (20x magnification). **(G)** Location of the RV+ last order premotor neurons after 55h of infection. RV+ neurons were represented by colored dots in the different segments of the cervical spinal cord for each preparation (prep #1 cian, prep#2 purple, prep#3 blue). Ipsi: ipsilateral side of the RV injection. **(H)** Plot of the number of RV+ neurons (# of neurons, x axis) in ipsilateral and contralateral sides and location along (distance, millimeters, y axis) the rostro-caudal cervical (C2-C8) segments for each preparation (prep #1 cian, prep#2 purple, prep#3 blue). **(I)** Number of RV+ motoneurons (MN, x axis) in the abducens and corresponding number of cervical last-order premotor neurons (pre-MN) for each preparation (prep #1 cian, prep#2 purple, prep#3 blue). **(J)** Mean +/-SD (black dot, 6.83±3.33) of the infection ratio indicating the average of last-order premotor neurons labelled per abducens motoneuron infected. **(K)** Scheme of the distribution of all the RV+ cervical neurons, shown in (G), *per* lamina of the cervical spinal cord.

### Coupling between forelimb and eye movements during sustained trot-like locomotion

To determine if locomotor-induced eye movements occur in the absence of visual and vestibular sensory signals, eye movements were recorded in the dark during head-fixed treadmill locomotion in decerebrated adult mice (Figure S4A). Such preparations enable the study of the spinal cord output and sensorimotor integration in the absence of descending supra-spinal motor commands^7^. Binocular nystagmic-like eye movements were observed during treadmill-evoked bouts of 10-40s sustained locomotion (Figure 4A, expanded section in Figure 4B). This locomotor-induced oculomotor behavior consisted in an alternation of short ocular movement (quick-phase like) in one direction, followed by longer eye movements (slow phase-like) of comparable amplitude in the opposite direction (Figure 4A-C). Eye movements were mostly restricted to the horizontal plane (compare vertical, in light colors, and horizontal components, in dark colors, in Figure 4B and 4C) and their orientation remained constant (Figure 4D) throughout the recordings. Left and right eye movements were always conjugated, with similar directionality for both eyes, and appeared well coordinated (see quick and slow eye movement phases on mean cycles of panel C; horizontal binocular coordination on oculogram in Figure 4E).

**Figure 4.**
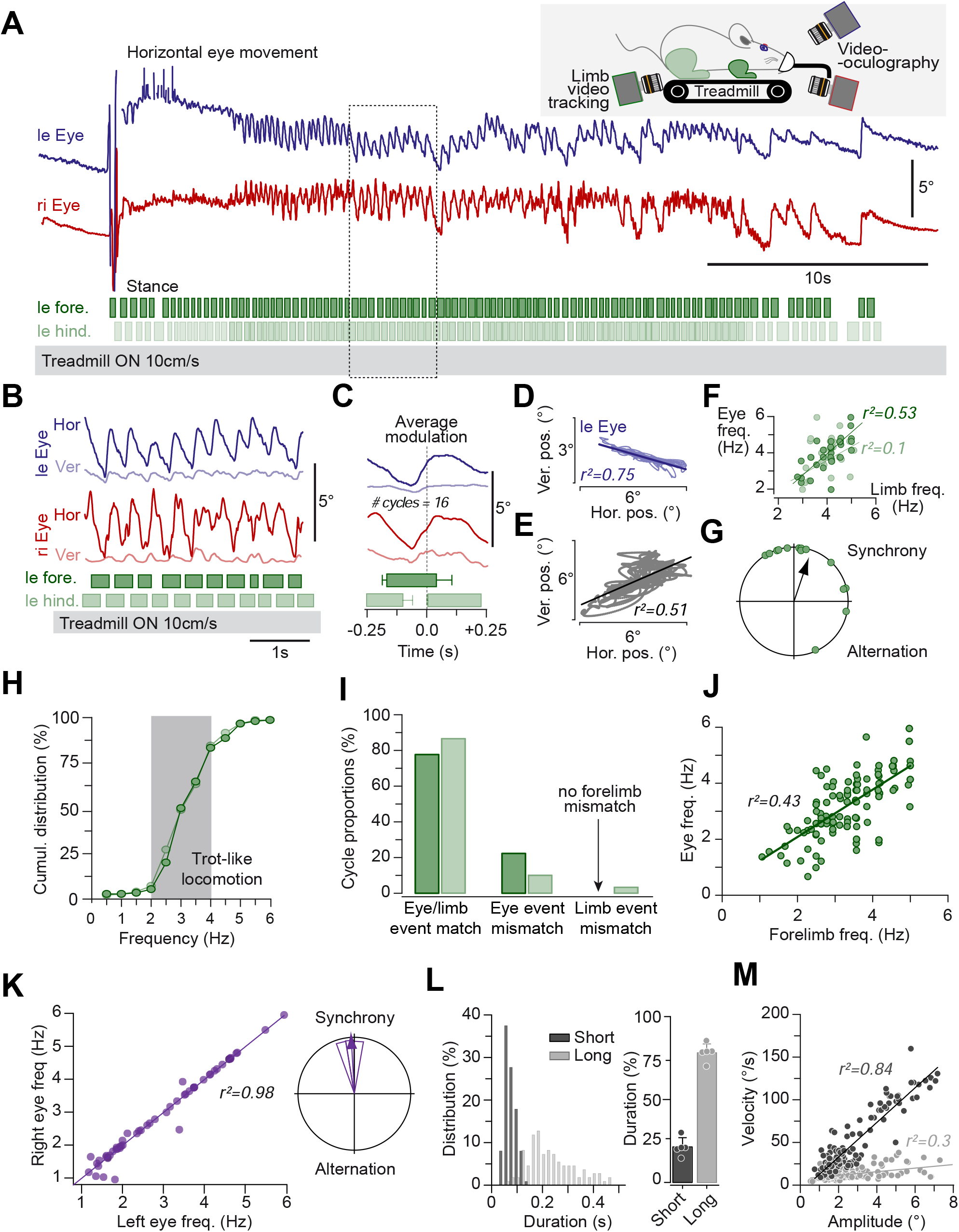
Locomotion-induced eye movements in adult mice. **(A, B, C)** Example of eye movements observed during an episode of treadmill-induced locomotion. Left (Leye, blue trace) and right (Reye, red trace) horizontal movements are observed after the onset of locomotion (dark green rectangles-left forelimb; light green rectangles-left hindlimb stance phases; gaps-swing). Upper right inset: depiction of the set up used for recording eye movements in decerebrated mice. Each eye had a video oculography camera recording its movements and an infra-red camera tracked the limb movement while mice ran on a motorized treadmill. **(B)** Segment (dotted rectangle, panel A) showing the horizontal (dark blue and red traces) and vertical (light blue and red traces) components of the eye movements and the corresponding locomotor cycles. **(C)** Average modulation of the eye movements and stance phases corresponding to panel B. For horizontal traces, right is up. **(D, E, F, G)** Analysis of the eye movements in panel **(B). (D)** Oculogram showing the constant orientation of the movement of the left eye, Hor.pos. vs Ver.pos (R=-0,8699, Spearman test, p=0,031; r² =0.75). Combined horizontal components of the right **(E)**, Hor.pos. vs Ver.pos, R=0,7152, Spearman test, p=0,47; r²=0.51) and left eyes showing comparable amplitude and synchronized movements. **(F)** Relation between the instantaneous frequency of the forelimb (dark green dots) or the hindlimb (light green dots), and the left horizontal eye movements () (Right eye vs left forelimb, R= 0,7248, Spearman test, *p <0,001;* r² =0.5253; Right eye vs left hindlimb, R= 0,3195, Spearman test, *p =0,0852;* r² =0.1021). **(G)** Polar plot of the phase coupling between eye and forelimb peak movement (µ= 18,299°±2,546; r= 0,718). **(H, I, J)** Relation between limbs and eye movements. **(H)** Cumulative distribution of the locomotor frequency. Eye movements were observed when locomotion frequency reached frequencies of 2-4 Hz, corresponding to trot-like gait (gray interval, n=6 mice). **(I)** Proportions of the locomotor cycle showing eye and/or limb movements (left forelimb-dark green; left hindlimb-light green). **J)** Instantaneous frequency (Hz) of the eye and forelimb movements (Forelimb vs eye instantaneous frequency, R= 0,6569, Pearson test, *p <0,0001;* r² =0,4315; n=6 mice). **(K, L, M)** Quantification of eye movements during locomotion. **(K)** Instantaneous frequency of the left and right eye movements. The conjugation of the eye movements is reflected in the high linear correlation (Left vs right eye instantaneous frequency, R= 0,9873, Pearson test, *p <0,0001;* r² =0,9748) between left and right eye frequency and the synchrony between them (right panel, polar plot; µ= 355,641°±7,53; r= 0,991). **(L)** The distribution of the duration of eye movements shows that short phases of the eye movements only last up to 200 milliseconds and are mainly distributed around 100 milliseconds while the long phase of the eye movements are scattered and distributed up to 400 milliseconds. Right panel shows the relative duration of brief and long eye movements during each cycle. Short eye movements (dark gray) represent 20% of the cycle; long eye movements (light gray) 80% of the cycles. **(M)** Main sequence of the eye movements showing the amplitude-velocity relationship of short (dark gray; Amplitude vs velocity of short component of eye movements, R= 0,9169, Spearman test, *p=*0,00366; r² =0,8395) and long (light gray; Amplitude vs velocity of long component of eye movements, R= 0,5412, Spearman test, *p<0,00001;* r² =0,2929) components of the eye movements.

**Table 1.**
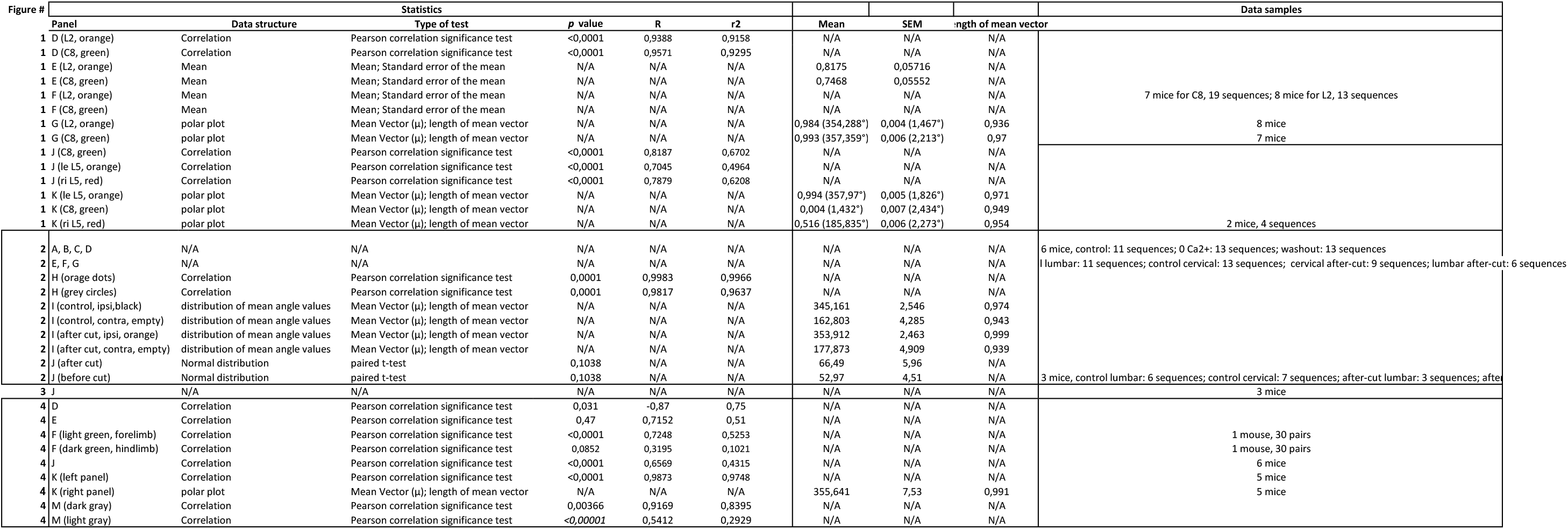

To determine the temporal relationship between locomotor and ocular movements, the occurrence of both limbs and eye events were compared. The frequency of the oculomotor cycles increased with the frequency of the locomotor cycles (Figure 4F). Moreover, eye and forelimb movements appeared in most instances synchronized (Figure 4G). All recordings shared two main features: first, consistent eye movements only occurred once a trot-like locomotion frequency (2-4Hz)^12^ was attained (Figure 4A and 4H). Second, the quantification of the event occurrence comparing eye, forelimb and hindlimb movements showed a match between eye and limb movements in about 80% of the cycles (Figure 4I, left histogram), and an eye mismatch (limb movement, no eye movement; Figure 4I, middle histogram) in about 20% of the cycles. Notably, locomotor-induced eye movements never occurred in the absence of forelimb movements (Figure 4I, right histogram). The frequency of the eye movement was closely correlated with the frequency of the forelimb movement (Figure 4J slope = 0.7882; *r*^*²*^=0.43). Left and right eye movements were tightly correlated and synchronized (*r*^*2*^=0,98; Figure 4K; phase in right panel). In all recordings, eye movements consisted in nystagmic-like movements composed of alternated short (0,066±0,022) and long (0,223±0,095) duration phases (Figure 4L, left panel) lasting 20% and 80% of the oculomotor cycles, respectively (Figure 4L, right panel). Both components had similar amplitudes in range 1-8° (Figure 4M, quick phases -black-, slow phases -gray-, and Figure S4E), leading to different velocities (amplitude/velocity relation on Panel M; slopes of 20.27 and 2.09 for short and long eye movements, respectively; Figure S4F). Overall, eye movements were observed during >80% of trot-like locomotor cycles and closely matched forelimb movements. Locomotor-induced synchronized binocular movements consisted in the alternation of short-duration rapid quick phase-like movements followed by longer duration slow phase-like eye movements in the opposite direction. The results obtained in decerebrated preparations thus confirm the presence of a spino-extraocular motor coupling based on an efference copy from the locomotor cervical CPG described in newborn mice (Figure 1 and 2). Additionally, they demonstrate that the persistence of this functional short-latency pathway in adults (Figure 3) ensures a tight coupling between forelimb locomotor movements and eye movements during sustained trot-like locomotion (Figure 4).

## Discussion

### Conservation of locomotor-induced spino-extraocular motor coupling through vertebrate evolution

Locomotor-induced spino-extraocular motor coupling was initially described in Xenopus frog^2– 6^. During swimming, both larvae and adult frogs exhibited a CPG efference copy-based extraocular bursting activity tightly coupled to the spinal motor rhythmic discharge. This spino-extraocular motor coordination adapted through metamorphosis to produce appropriate horizontal eye movements that counteracted either tail (larva) or limb (adults) movements during strong propulsive swimming^4–6^. The efference copy mechanism was supported by an ascending spinal-brainstem neural pathway connecting spinal neurons to extraocular motoneurons, crossed in larvae and bilateral in adult xenopus^3,5^. Using newborn and adult mice preparations, the present study reports a locomotor-induced spino-extraocular motor functional coupling that exhibits common features with frogs. First, in both species this coupling is supported by a direct monosynaptic pathway that persists during mouse ontogeny, as previously reported in xenopus. Second, the bilateral distribution of abducens-projecting cervical neurons revealed in mice by RV tracing suggests a comparable organization of the spinal-brainstem pathway between adult frog and mouse, putatively common to tetrapod locomotor systems. Third, the locomotor-induced ocular signal, recorded either *ex vivo* or in decerebrated animals, was only observed during sustained and vigorous locomotor activity, corresponding to running behavior, a terrestrial equivalent for propulsive undulatory swimming. *Ergo*, our findings, combined with previous studies in Xenopus, strongly support the hypothesis that the CPG efference copy-based ocular signal is a conserved mechanism in amphibians and mammals, and putatively in all vertebrates.

This conserved locomotor-induced ocular behavior is in accordance with the absolute necessity to stabilize the visual field during motion, by performing rapid counteracting eye adjustments in all vertebrates, including humans^13^. There is considerable behavioral evidence for the contribution of locomotor-related spinal signals during gaze stabilization in several mammalian species. In cat^14,15^ and in humans^16^, locomotor signals are integrated to favor vertical eye movements by projecting gaze towards the direction of locomotion and canceling VOR during walking or running on a treadmill. In monkeys performing circular running in the dark, compensatory horizontal eye movements are favored by non-vestibular velocity signals^17^. Here, we report new evidence demonstrating the existence of the spino-extraocular motor command in mice that could constitute a neural mechanism sustaining the coupling of locomotion and eye movements, as suspected of other mammalian species.

### Adaptation of the CPG efference copy-based mechanism through the vertebrate lineages

Gaze control largely results from the sensory-motor transformations which primarily depend on the integration of visual and vestibular inputs (optokinetic and vestibulo-ocular reflex)^18^. Interestingly, no locomotor-induced ocular movements were observed in head-fixed alert mice during sustained treadmill locomotion in the dark (Figure S4), suggesting a sensory-dependent supraspinal control of the CPG efference copy signal in natural conditions, which is suppressed in decerebrated mice. Recent studies in mice reported that the vestibulo-ocular reflex accounts for a dominant part of the conjugated gaze stabilization observed during head-free locomotion ^19–21^. VOR-based eye movements were found to reach higher dynamics, compensating naturalistic head movements interspersed with non-conjugated (vergence) and shifting movements relocating gaze during directed head turn^21,22^. The spino-extraocular coupling observed in our study consisted in a symmetrical and conjugated ocular pattern. This suggests that the locomotor efference copy signal might contribute to a set of conjugated, eye movements. Whether this mechanism is participating in compensatory and/or anticipatory eye movements during quadrupedal running behaviour remains to be determined. Even though the spino-extraocular motor command is the predominant way to stabilize gaze during swimming in frogs, this mechanism might complement the visuo-vestibular based reflexes during high dynamic locomotor patterns in mice. All in all, the evidence suggests an adaptation of the CPG efference copy-based mechanism through the vertebrate lineages in relation with changes in sensory capacities and locomotor repertoire during evolution.

## Figure Legends

**Supplementary Figure 1.**
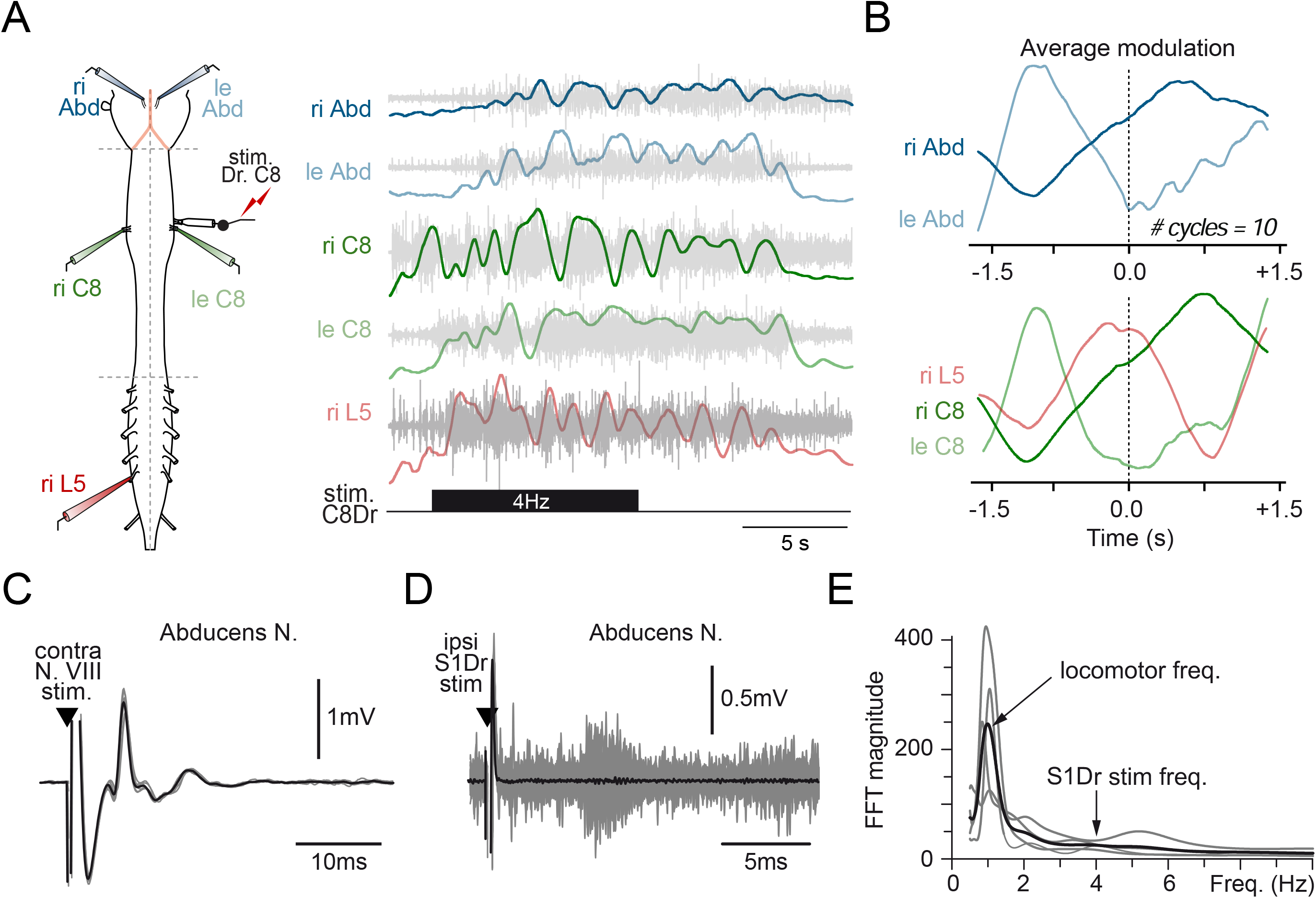
**(A)** Fictive locomotion episodes evoked by cervical CPG stimulation. Schematic of the neonatal mouse brainstem and spinal cord preparation with stimulated and recorded nerve branches (left side). Extracellular nerve recordings (right side, raw traces in light gray, integrated traces in color) of the left (le, light blue) and right (ri, dark blue) abducens nerves (Abd.), the left and right 8th cervical roots (le C8, light green; ri C8, dark green) and the right 5th lumbar ventral root (ri L5, red). Discharges recorded during an episode of fictive locomotion evoked by the electrical stimulation of the C8 dorsal root (stim. S1Dr) with a 4Hz pulse train (black rectangle). **(B)** Average cyclic modulation of the discharge activity (integrated trace) from the motor nerves shown in (A) over 10 consecutive locomotor cycles. The ri C8 trace (dark green) was used as the reference to determine locomotor cycles **(C)** VIIIth nerve-evoked (black arrowhead) abducens motor responses (individual traces in gray; mean in black) occur with a typical disynaptic latency compatible with the activation of the direct VOR pathway. **(D)** Superimposed abducens activity in response to electrical pulse (black arrowhead) applied on S1Dr (gray traces) and average activity (black trace). No reproducible S1Dr-related activity was observed following locomotor-evoked abducens discharge. **(E)** Frequency periodograms obtained from FFT (Fast Fourier Transform) analysis of the abducens nerve discharge during the S1Dr evoked fictive locomotion episodes (grey traces represent a single locomotor sequence, the black trace is the average from the 4 grey traces). Periodograms revealed only one magnitude peak at the locomotor frequency (locomotor freq., about 1Hz) and did not shown any peak at the S1Dr stimulation frequency (S1Dr freq., 4Hz), demonstrating the absence of spinal afference-evoked direct response in the abducens discharge.

**Supplementary Figure 3.**
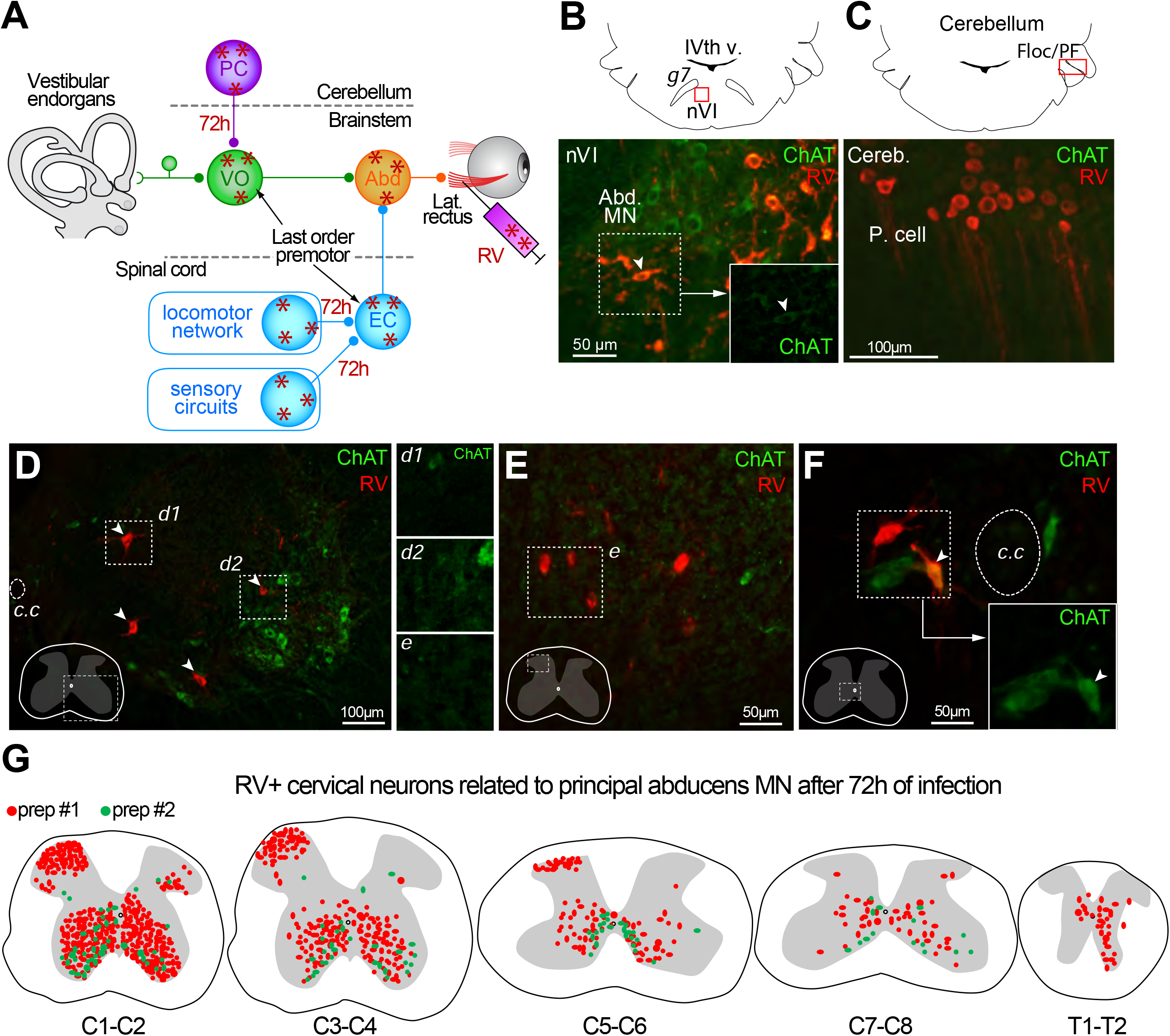
**(A)** Depiction of the disynaptic infection obtained 70h after inoculation in the lateral rectus muscle. RV+ neurons are represented with red asterisks. Abd, abducens nuclei; PC, Purkinje cells; VOR, Vestibulo-ocular central neurons; EC, efferent copy spinal neuron. Sp, spinal neuron X-RV+ interneurons along the spinal cord after a disynaptic infection. **(B)** Representation (top panel) of the location (red box) of the RV+ neurons in the abducens nucleus (nVI) and its landmarks: geniculum of the facial nerve (g7, 7n) and IVth v, 4th ventricle. The bottom panel shows a fluorescence microscopy image (20x) of an RV+/ChAT+ neuron (RV, red) and abducens motoneurons (Abd. MN, ChAT, green) 55h after the infection. **(C)** Representation (top panel) of the cerebellar location (red box, Floc/PF: Flocculus/Paraflocculus) of the RV+ Purkinje cells (P. cell) shown in the fluorescence microscopy image (bottom panel, 20x magnification) 70h after the RV infection. **(D, E, F)** Fluorescence microscopy images of different cervical spinal cord slices and the ventral (**D**) (10x magnification), dorsal (**E**) horns (20x magnification) and around the central canal (c.c) (**F**) (20x magnification) location of RV+ neurons. The RV+ neurons found in the ventral (**d1, d2** insets) and dorsal (**e** inset) horns were not ChAT+. However, some neurons found around the central canal (**f**, inset) were simultaneously RV+ and ChAT+. **(G)** Scheme of the location of the RV+ neurons found in the different cervical (C1-C8) or thoracic (T1, T2) segments following the 70h-disynaptic protocol. Red and green dots correspond to 2 different preparations (prep#1 red; prep #2 green). Labeled neurons were found with a clear rostro-caudal gradient, in the ipsilateral and contralateral ventral horn, and in the ipsilateral dorsal horn of rostral cervical segments.

**Supplementary Figure 4.**
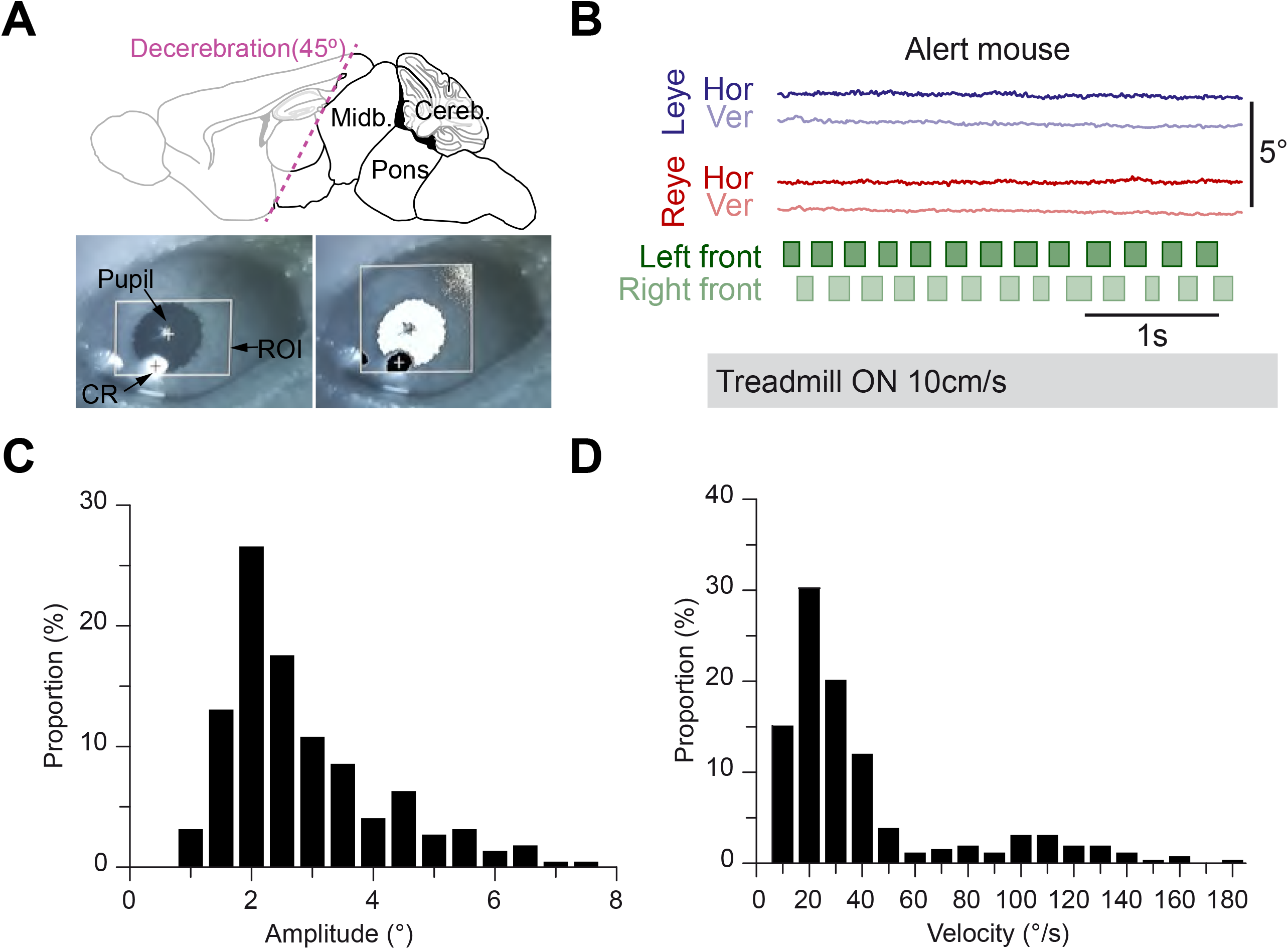
**(A)** Upper panel: scheme of the decerebration performed in adult mice. The 45° decerebration angle ensured a premammillary and precollicular decerebration; the structures outlined in black were preserved while the ones in gray were removed. Lower panel: picture of the right eye during a videoculography recording with region of interest (ROI) tracked; the pupil and corneal reflection (CR). The right image shows the same eye with the detection threshold applied. **(B)** Eye movement traces and locomotor cycles of an intact mouse (head-fixed but not decerebrated) running on a treadmill at 10 cm/s. Neither left (Leye; dark blue trace: horizontal, light blue trace: vertical) nor right (Reye; dark red trace: horizontal, light red trace: vertical) horizontal and/or vertical movements are observed after the onset of locomotion (dark green-left forelimb; light green-left hindlimb stance phases) Distribution of the Amplitude (**C**) and velocity (**D**) of the eye movements generated during treadmill locomotion in decerebrated preparations (9 mice).

## STAR Methods

### Key Resources Table

| REAGENT or RESOURCE | SOURCE | IDENTIFIER |
| --- | --- | --- |
| **Antibodies** |  |  |
| Anti-RV phosphoprotein | Coulon et al 2011 | N/A |
| Anti-Choline Acetyltransferase(ChAT) antibody | Merck | AB144P |
| Goat anti-Mouse IgG (H+L), Superclonal™ Recombinant Secondary Antibody, Alexa Fluor 546 | Thermo Fisher Scientific | Cat # A10036 |
| Donkey anti-Goat IgG (H+L) Cross-Adsorbed Secondary Antibody, Alexa Fluor 488 | Thermo Fisher Scientific | Cat# A11055 |
| **Bacterial and Virus Strains** |  |  |
| Challenge Virus Strain, CVS | Gaudin et al., 1992 | N/A |
| **Biological Samples** |  |  |
| x |  |  |
| **Chemicals, Peptides, and Recombinant Proteins** |  |  |
| x |  |  |
| **Critical Commercial Assays** |  |  |
| x |  |  |
| **Deposited Data** |  |  |
| Dataset_Barros_2021 | This study | Database: Mendeley<br>doi: 10.17632/h8nbswgf95.1 |
| **Experimental Models: Cell Lines** |  |  |
| x |  |  |
| **Experimental Models: Organisms/Strains** |  |  |
| x |  |  |
| **Oligonucleotides** |  |  |
| x |  |  |
| **Recombinant DNA** |  |  |
| x |  |  |
| **Software and Algorithms** |  |  |
| DeepLabCut | Mathis et al., 2018 | github.com/AlexEMG/ |
| DataView | W.J. Heitler, University of St.Andrews Scotland | https://www.st-andrews.ac.uk/~wjh/dataview/ |
| Spike2 | Cambridge Electronic Design | http://ced.co.uk/products/spk2/ |
| Prism | GraphPad | https://www.graphpad.com/scientific-software/prism/ |
| Oriana | Kovach Computing Services, United Kingdom | https://www.kovcomp.co.uk/oriana/ |
| Zeiss ZEN | Carl Zeiss | https://www.zeiss.com/microscopy/us/products/microscope-software/zen-lite.html |
| **Other** |  |  |
| Data acquisition interface | Cambridge Electronic Design | Power1401 MkII |
| Video-oculography system | ISCAN, Burlington, MA | ETL-200 |
| Infrared video camera | FLIR Systems, Inc. | Grasshopper3 |
| Motorized treadmill | Panlab, Harvard Apparatus | Item#76-0891 |
| Low-flow anesthesia system | Kent Scientific Corporation | SomnoSuite |
| AC amplifiers | A-M Systems, Carlsborg, US | Model 1700 |
| Stimulus isolator (A.M.P.I.) | Science Products, GmbH | ISO-Flex |
| Ventilator | Kent Scientific Corporation | PhysioSuite |

### Resource availability

#### Lead contact

Further information and requests for resources should be directed to M. Beraneck.

#### Materials availability

This study did not generate any new materials or reagents.

#### Data and code availability

The datasets generated during this study are available at “Dataset_Barros_2021”, Mendeley Data, V1, doi: 10.17632/h8nbswgf95.1

### Experimental model and subject details

Animals were used in accordance with the European Communities Council Directive 2010/63/EU. All efforts were made to minimize suffering and reduce the number of animals included in the study. All protocols were performed on C57/BL6J male mice. *Ex vivo* experiments were performed on neonate (2-3 days old; n= 23) following procedures specifically approved by the ethical committee for animal research of the University of Bordeaux. Experiments on adults (8–14 weeks) were approved by the ethical committee for animal research of the University of Aix-Marseille (n=6) and of the university of Paris (n=29).

### Ex vivo brainstem-spinal cord preparations

The dissection protocol followed as previously detailed by Kasumacic^10^. Briefly, mice were deeply anesthetized by 4% isoflurane inhalation until the loss of the nociceptive reflexes. After being placed in Sylgard-coated Petri dish, they were decerebrated and eviscerated while submerged in an ice-cold (1°-5°C) oxygenated (95% O_2_ and 5% CO_2_) artificial cerebrospinal fluid (aCSF) (in mM: 128 NaCl, 4.5 KCl, 2.5 CaCl2.2H2O, 1.0 MgSO_4_.7H_2_O, 1.2 NaH_2_PO_4_.H_2_O, 5 Hepes, 25 NaHCO_3_ and 11 D-glucose). In a dorsal view, a craniotomy was performed to expose and remove the cerebellum, and the abducens nerves were carefully cut so that they could be recorded. With a ventral approach, a full laminectomy was done to expose the spinal cord and all the dorsal and ventral roots of both cervical sides were cut so that the final brainstem-spinal cord preparation could be extracted. After removing the meninges, the preparation was fixed with the ventral side up and fixed to the Sylgard-coated dish with insect pins and bathed in oxygenated aCSF fluid at room temperature.

### Fictive locomotion protocols and recordings

Isolated brainstem-spinal cord preparations had the distal ends of abducens nerves as well as bilateral spinal dorsal motor roots (sacral, lumbar or cervical) simultaneously recorded (Model 1700 AC amplifiers, A-M Systems, Carlsborg, US) with fire-polished borosilicate glass suction recording electrodes filled with aCSF solution. The electrodes were connected to extracellular amplifiers and the analog signals recorded by an analog/digital interface (CED 1401; Cambridge Electronic Design, Cambridge, UK). Data was acquired with Spike2 software (Spike2, Cambridge Electronic Design) at a frequency of 1000Hz. Fictive locomotion was achieved in brainstem-spinal cord preparations either through electrical or pharmacological stimulation.

For electrical stimulation, the electrodes suctioning the dorsal roots were connected to a pulse generator; this allowed the delivery of pulse trains (4Hz for 10s, inter-stimulus interval of 0.25s and at least 5 minutes between pulse trains). Signals were amplified (x10000) by differential AC amplifiers, digitized at 10 KHz (CED 1401; Cambridge Electronic Design, Cambridge, UK), displayed and stored on computer with acquisition software (Spike 2, Cambridge Electronic Design) and finally analysed off-line with customized scripts. The described pulse trains were delivered at the lumbar or at the cervical levels while the abducens nerves were recorded.

To obtain fictive locomotion through pharmacological stimulation, the *ex vivo* brainstem-spinal cord preparations (n= 2 mice) were bathed in an aCSF infused with glutamatergic receptor agonists; namely NMDA (7,5 µM) and serotonin (5-HT; 15 µM). The recordings were performed as described above.

In a subset of experiments (n=6 mice), the motor coupling was disrupted using a split-bath configuration that allowed reversibly isolating the brainstem from the spinal cord. Isolation was obtained by partitioning the recording chamber with two custom-made plastic walls that traversed the preparation cervical and lumbar regions with spinal split-baths. The cervical spine was immersed in an aCSF 0 Ca2+ solution, in order to completely block transmission at that level, while lumbar root stimulation was performed. To isolate the cervical from the rest of the preparation, a section was made at the thoracic spinal cord and the cervical was then stimulated while the abducens nerves were recorded.

### Methods for tracing experiments

Rabies virus (RV) manipulation was done by vaccinated experimenters in a Biosafety level 2 facility. The virus (Challenge Virus Strain, CVS) was produced and concentrated as previously described^23^. Aliquots of the virus were stocked at -80°C and thawed before use. Mice were anesthetized by an i.p. injection of ketamine (Imalgene, 60 mg/kg) and xylazine (Rompun, 10 mg/kg) and deep anesthesia was confirmed by lack of response to interdigital pinching. The left extraocular muscles were exposed through an incision on the skin covering the mouse’s eye, surrounding fatty tissue was removed and absorbable hemostatic gelatin sponges were used to curtail bleeding (Spongostan, Ethicon). After its identification, the lateral rectus muscle (LR) was isolated using small hooks and an injector cannula (gauge 33), linked to a 10µL Hamilton syringe, was inserted. 1µL of the RV was then slowly injected and the needle was left in place for an extra minute to prevent leaking^24,25^. Finally, the skin was closed using non-absorbable monofilament suture and mice were put back on their cages under a red light for post-surgery care. Labelling times were narrowed down to obtain RV-infected spinal cord neurons with monosynaptic connections to the abducens nucleus. A 70h labelling window displayed RV+ neurons in the cerebellum (Purkinje cells; Figure S3C), confirming disynaptic connections to that delay. When the inoculation time was lowered to 55h the RV+ neurons were restricted to the brainstem and cervical spinal cord, with no infection on the cerebellum or any adjacent areas.

### Immunochemistry

After 55 or 67hh of RV infection, animals were deeply anesthetized with ketamine (Imalgene, 60 mg/kg) and xylazine (Rompun, 10 mg/kg) and perfused intracardially with 0.1M phosphate buffer saline (PBS) followed by 4% paraformaldehyde (PFA). Immediately after, the brain was dissected coronally from the midbrain to the medulla oblongata and the spinal cord was removed in a single block. Tissue fixation was let for 4h at 4°C in 4% PFA, subsequently cryoprotected in 20% sucrose during at least 24h and freeze embedded at -80°C in Tissue Tek (Sakura, Zoeterwoude, The Netherlands). Transverse sections (30 µm thick) of the brain and cervical, thoracic and lumbar spinal cord were serially cut in cryostat (Microm, Heidelberg, Germany) and collected onto poly-L-lysine-coated slides. These sections were first incubated to avoid nonspecific binding for 1h at room temperature in blocking solution (PBS, Triton X-100, 0,2% bovine-serum albumin and normal donkey serum) and then left at 4°C for 48h with the anti-RV phosphoprotein^25^ and anti-choline acetyltransferase (ChAT) diluted 1:100 in the blocking solution. After being rinsed thrice in PBS, sections were incubated at room temperature with the secondary antibodies Alexa546 goat anti-mouse (Thermo Fisher Scientific) and Alexa488 donkey anti-goat (Thermo Fisher Scientific) at a 1:200 dilution for 2h. Finally, sections were rinsed two times with PBS and once with tap water before being mounted (Immu-Mount, Fisher Scientific) under coverslips.

To analyze the distribution of RV+ neurons, the obtained sections were visualized under an upright light microscope (Axioscope, Carl Zeiss, Germany). Images of each section were taken using a CCD camera (AxioCam MR3, Carl Zeiss). These images were afterwards treated in ZEN (Carl Zeiss) imaging software where brainstem RV^+^ /ChAT^+^ motoneurons and spinal cord RV^+^ neurons were counted on each section and plotted from one out of two (60µm apart). Only neurons with visibly infected nuclei, determined by the presence of Negri bodies, were considered.

The success of the viral infection was verified by the presence of double-positive RV+ and ChAT+ neurons on the abducens nucleus (Figure 3B). Abducens motoneurons simultaneously RV+ and ChAT+ were found in all mice (n=3), ipsilateral to the injection site (left lateral rectus muscle), while motoneurons innervating other eye muscles (Figure 3B) were never marked, confirming the specificity of the infection.

### Procedure for decerebration

Procedures for decerebration were previously described by Meehan and colleagues^7^. Briefly, mice were deeply anesthetized with isoflurane (2–3% in 100% O2) in an induction chamber, and then transferred to a heating pad on the surgery table with a nose cone to maintain anesthesia (1.5–2% in 100% O2). A rectal thermometer (Physiosuite, Kent Scientific) was used to monitor body temperature throughout the surgery as well as a pulse oximeter (MouseSTAT, Kent Scientific) to monitor O_2_ saturation. Mice were then artificially ventilated via tracheostomy by blunt dissection; first the sternohyoid muscle was exposed, then the upper half of the fibrous membrane between two cartilages was cut and finally, the air delivery mode was quickly switched to ventilator mode (SomnoSuite, Kent Scientific) and a plastic endotracheal tube was inserted into the trachea. After confirming that the pCO2 levels were acceptable (>2.5%), the isoflurane was lowered to 1-1,2%. To avoid excessive bleeding during the decerebration, the left and right carotid arteries were bluntly isolated, ligatured and cauterized using an electric cauterizer (Change-a-tip cauteries, Bovie). The initial midline incision was sutured and, to avoid dehydration, 0.3 ml of sterile lactated Ringer’s solution (Braun Medical) were injected subcutaneously. The mouse was placed in a stereotaxic frame and a superficial cut was made across the skin above the midline of the skull. Using a micro-drill (Foredom, David Kopf Instruments), the skull was scored and the parietal bones removed. Decerebration was performed at the confluence of the sagittal and transverse sinus, perpendicular to the midline and at a 40±5° angle (Figure S4A). A surgical scalpel (no.10 blade; Fine Science Tools) was used to perform a swift and gentle slicing motion. The structures rostral to the incision were fully removed to obtain a premammillary and precolliculary decerebration. Finally, a subcutaneous injection of Ringer’s lactate (0.3 ml) was applied, and the isoflurane gas anesthesia was decreased to zero. Complete anesthesia withdrawal and muscular tonus occurred 10-20 minutes after isoflurane removal; PCO_2_ levels rose consequently.

### Videoculography and locomotion recordings and analysis

After the decerebration, mice were tightly secured using ear bars and a custom-made mouthpiece onto the motorized treadmill (Panlab, Harvard Apparatus). All recordings were performed in complete darkness. The locomotion was recorded using an infrared video camera (100Hz; Grasshopper3, FLIR Systems, Inc) fixed perpendicular to the left side of the body. To perform binocular video-oculography, the eyes of the mouse were illuminated with infrared emitters attached to 2 CCD cameras (120Hz; ISCAN, Burlington, MA) that were placed symmetrically on each side of the treadmill. Special care was provided to prevent eye dryness by regularly re-applying ophthalmic gel, to ensure that at least one eye could be tracked online. Eye video signals were processed online (ETL-200, ISCAN, Burlington, MA), sampled at 1kHz (CED 1401; Cambridge Electronic Design, Cambridge, UK) and recorded with Spike2 software. Online tracking of the eye movements with set-up fixed cameras allowed verifying the absence of head movements during the recordings. The video-oculography and locomotion cameras were synchronized using trigger signals generated using spike2 software. Locomotor cycles were tracked from the acquired videos using a markerless pose estimation (DeepLabCut)^26^. Eye movements and locomotion were analyzed offline using DataView software (W.J. Heitler, University of St.Andrews Scotland).

Overall, eye movements coupled with locomotion were observed on n=26 preparations. Because the spino-extraocular coupling critically depends on the locomotion performed (see results), only preparations which exhibited bouts of sustained (>30s) locomotion at frequency >1Hz were retained (n=9, including 5 with binocular recordings) for final reporting.

### Quantification and statistical analysis

A table featuring all the numerical values obtained from the statistical tests performed is given in the Supplementary information. Prism (GraphPad) was used for statistical analysis.

For paired t-tests normality was first evaluated using the D’Agostino & Pearson normality test and Shapiro-Wilk in the case of smaller unpaired two-tailed samples. When testing locomotion frequency and latency as well as the integrated electrophysiological signals, differences between two results were obtained using the unpaired two-tailed Mann–Whitney *U*-test and Kolmogorov–Smirnov test to compare distributions. To compare several values, the non-parametric Kruskal–Wallis test was processed with a Dunn’s multiple comparisons test. To evaluate the correlations between the frequencies of discharge of the locomotor nerves and the extraocular nerves, regression tests (R) and linear regression (r²) were performed. Regression results were expressed as (R, r²). Circular data was analysed with Oriana 4.02 (Kovach Computing Services, UK) and the phase and strength of coupling were indicated by their mean vector (µ) and its length (r), respectively for non-uniform distributions values (tested with the Rayleigh’s uniformity test, p).

The results are expressed as the mean ± SEM and *p* values threshold determined as ^∗^*p* < 0.05; ^∗∗^*p* < 0.01; ^∗∗∗^*p* < 0.001; ^∗∗∗∗^*p* < 0.0001; ns: non-significant.

## Acknowledgements

The authors thank Patrice Jegouzo for his technical help designing the set up for the semi-intact experiments, Patrice Coulon for his input on the rabies virus tracing and the Animotion platform (INCIA) for the expertise in the locomotion analysis.

FFB, CT, MT & MB were supported by the Centre National d’Etudes Spatiales, the Centre National de la Recherche Scientifique and the Université de Paris, and by the Agence Nationale de la Recherche (ANR-15-CE32-0007-02).

This study contributes to the IdEx Université de Paris ANR-18-IDEX-0001.

This work has benefited from the support and expertise of the animal facility of BioMedTech Facilities at Université de Paris (Institut National de la Santé et de la Recherche Médicale Unité S36/Unité Mixte de Service 2009).

## Author contribution

Conceptualization: FB; FML, DC, MB. Methodology: FB; MM, JBC, MT, HB, FML, MB. Software: JBC, MM. Formal analysis: FB; JBC, MT, CT; MM, FML, MB. Investigation : FB; CT, MM, HB. Writing original manuscript: FB; JBC, FML, MB. Writing review and editing: FB; MM, JBC, MT, HB, FML, MB. Visualization: FB; FML; MB. Supervision: FML; MB. Project administration: DC, FML, MB. Funding acquisition: DC, FML; MB.

## Supplemental information

